# Lipidomics and colistin resistance in non-human isolates of *Acinetobacter seifertii*

**DOI:** 10.1101/2024.05.22.595387

**Authors:** Ellen M. E. Sykes, Valeria Mateo-Estrada, Anna Muzaleva, George Zhanel, Jeremy Dettman, Julie Chapados, Suzanne Gerdis, Izhar U. H. Khan, Santiago Castillo-Ramírez, Ayush Kumar

## Abstract

*Acinetobacter baumannii* is most well known for its role as a human pathogen and as a member of the *Acinetobacter calcoaceticus-baumannii* (ACB) complex. However, lesser characterised members of the ACB complex, have also been implicated in hospital-acquired infections. Once mainly considered opportunistic pathogens, many *A. baumannii* and non-*baumannii* strains are being isolated from agricultural, water and food sources. The surveillance and tracking of *Acinetobacter* spp. have been recently suggested to be part of the One Health consortium, to understand and prevent the spread of antimicrobial resistance. Here, we isolated four *Acinetobacter* strains from tank milk in Bogor, Indonesia and using ANI and dDDH techniques have identified them as *Acinetobacter seifertii*. MLST methods assigned these *A. seifertii* strains to a novel Sequence Types (ST), highlighting the diversity not only within the ACB complex but also in non-human *Acinetobacter* spp. These four *A. seifertii* strains are colistin-resistant and while they do not harbour any known mechanism of colistin resistance, they do share amino acid substitutions in regulatory proteins, AdeS, PmrAB, H-NS, and the membrane associated proteins, LpxACD, MlaD, PldA, LpsB and EptA that may contribute to this phenotype. Furthermore, down-regulation of the RND efflux pump AdeAB, may also be a key factor in colistin resistance in these non-human *A. seifertii* strains. Lipidomics revealed an acyl-homoserine lactone (AHL) molecule, and lyso-phosphatidylethanolamine (lyso-PE) in significant abundance compared to colistin-sensitive *A. baumannii* ATCC17978 revealing lipidomic differences between species. Finally, these four tank milk *A. seifertii* strains are avirulent in an insect model of virulence. It is possible that *A. seifertii* strains are intrinsically resistant to colistin and require further study. By investigating these less understood *Acinetobacter* spp. from non-human sources, our study supports the One Health approach to combatting antibiotic resistance.

## Introduction

Members of the *Acinetobacter* genus have only recently been investigated for their clinical relevance. The development of the term “ACB complex” species includes *Acinetobacter pitti, A. nosocomialis, A. calcoaceticus,* more recently, *A. seifertii* and *A. lactucae* (1) and the most widely reported to be associated with hospital-acquired infections *Acinetobacter baumannii.* The World Health Organization has classified carbapenem-resistant *A. baumannii* as a priority one pathogen for which more research and development is required (2). Study of this pathogen and its relatives is critical because upwards of 70% of *A. baumannii* strains are multi-drug resistant (MDR) and some are even resistant to the last resort antibiotics carbapenems and colistin (3).

*Acinetobacter seifertii* was first classified as a separate species within the ACB complex in 2015 (4) and has more recently been isolated from clinical settings in Japan (5), Brazil (6) and Taiwan (7). Some clinical (6, 8, 9) and non-clinical (9) isolates displayed resistance to the last resort antibiotic, colistin, while others did not (10). Compared to other *Acinetobacter* spp. very little is understood regarding the *A. seifertii* population structure, global distribution, isolation sources and antibiotic resistance and virulence potential.

Although focus on *A. baumannii* and other ACB complex members has mainly been clinical, there is mounting evidence that investigation into non-human strains is essential. Our understanding of *Acinetobacter* lineages and antibiotic resistance lies beyond clinical infections and must include study of strains of environmental origin. In short, a One Health approach is required (11). By exploring the diversity of non-human *Acinetobacter* spp., further insight can be gathered into their evolution including novel STs, and antibiotic resistance genes, (12) as well as their potential to cause serious consequences for human health (13). In fact, emerging *Acinetobacter* spp, such as *A. junii* (14) and *A. berenziniae* (15) are being classified as One Health pathogens. Transmission of antibiotic resistance genes and their mobile genetic elements from non-human strains to concerning STs further supports a One Health approach to *Acinetobacter* research (16).

Colistin is a glycopeptide antibiotic, which was primarily used in hospitals to treat hospital-acquired infections bacterial infections. Due to toxicity, its use declined but has recently increased especially in agricultural practises, mainly due to the requirement for a last resort antibiotic (17). The use of colistin in livestock and veterinary medicine remains high, as a strategy for infection prevention and growth promotion among animals (18). The emergence of many MDR pathogens as the cause of human infections requires the administration of less-used and less-important in human medicine antibiotics, such as colistin, in order to successfully treat resistant human infections. The increase in clinical colistin use has also brought on an increase in colistin-resistant bacteria and is marked by an increased prevalence of the *mcr-1* (mobile colistin resistance) gene (19). Globally, the use of antimicrobials in all sectors is highest in countries such as China, USA, Brazil, Germany and Indonesia (20).

Colistin resistance in *A. baumannii* has been primarily characterized in clinical and reference strains and is associated with the modulation of membrane homeostasis. This includes modification or loss of the lipooligosaccharide (LOS), including the target of colistin, lipid A. Modification of lipid A, is a common colistin resistance mechanism in many bacterial species (21, 22). In *A. baumannii,* the addition of phosphoethanolamine modifies the charge of lipid A, thereby disrupting the cationic interaction of colistin and the negatively charged lipid A preventing membrane disruption. The two component system PmrAB, along with the phosphoethanolamine transferase, PmrC, play a role in regulating these lipid A modifications (23). Additionally, mutations in the global regulator H-NS cause upregulation of the *pmrC* homolog, *eptA*, also resulting in the decoration of lipid A and thus colistin resistance (24). Finally, the *mcr* gene and its derivatives are also known to add phosphoethanolamine to lipid A, and *mcr-1* (25)*, mcr-2*, *mcr-3* (26) and *mcr-4* (27) have all been detected in *A. baumannii*.

Complete loss of LOS leading to colistin resistance, is facilitated by the presence of a variety of insertion elements and mutations in membrane-associated biosynthetic pathways. The insertion of IS*Aba*11 in the lipid A biosynthetic genes *lpxC* and *lpxA* (28) as well as Single Nucleotide Polymorphisms (SNPs) and deletions in the same pathway resulted in the absence of LOS (29). Furthermore, the insertion of IS*Ajo*2 has been found to interrupt both *lpxA* and *mlaD* which encode membrane anchored periplasmic protein involved in the transport of glycerophospholipids. IS*Aba*13 has been identified interrupting *pldA* and has been linked with colistin resistance (30). The mechanism(s) of colistin resistance in non-*baumannii* species of *Acinetobacter* has not been studied in detail but due to phylogenetic relatedness, it is likely that similar mechanisms of colistin resistance are plausible in these species such as *A. seifertii*. Here we begin to explore colistin resistance in non-clinical isolates of this ACB complex member.

## Results

### Strains isolated from tank milk in Indonesia were identified as *A. seifertii*, represent novel sequence types

*Acinetobacter seifertii* strains, AB348-IK22, AB359-IK23, AB349-IK24, and AB350-IK25, were isolated from tank milk in Bogor, Indonesia. Their identity was confirmed as *A. seifertii* using the Average Nucleotide Identity based on MUMmer (ANIm) and digital DNA:DNA hybridization (dDDH) analysis. Table 1 reports these values in a matrix format. AB348-IK22 had an ANI value of 97.1% compared to *A. seifertii* UBA3035 (Accession: GCA_002367535.1) and 97.4% relative to *A. seifertii* DSM102854 (NIPH 973) (Accession: GCF_000368065.1). In fact, three tank milk *A. seifertii* strains (AB359-IK23, AB349-IK24 and AB350-IK25) had the same ANIm and similar dDDH values as AB348-IK22 (Table 1). Further, comparing ANIm values of all strains to the ATCC17978 (NZ_CP018664.1) *A. baumannii* strain, they were all below the 95% cutoff for species identification (31), indicating that these were not *A. baumannii* strains. Values for dDDH confirm ANI species identification. AB348-IK22 compared to *A. seifertii* UBA3035 and *A. seifertii* DSM102854 results in a dDDH value of 75.2 % (Table 1).

**Table 1:** Average nucleotide identity (ANI) and digital DNA:DNA Hybridization (dDDH) comparison for identification of AB348-IK22, AB359-IK23, AB349-IK24 and AB350-IK25. The ANI value for each strain is calculated in JSpecies and then a matrix generated. The cutoff for a species match for ANI is above 95% and above 75% for dDDH. AB348-IK22, AB359-IK23, AB349-IK24 and AB350-IK25 compared to the type strains UBA3035 and DSM102854 (NIPH 973) are all above ANI values of 97% and dDDH values of 75% indicating that they are in fact *A. seifertii*. The type strain of *A. baumannii* ATCC17978 is used as an out of species comparator.

|  | ANIm |  |  |  |  |  |  |
| --- | --- | --- | --- | --- | --- | --- | --- |
| Strain | AB348-IK22 | AB359-IK23 | AB349-IK24 | AB350-IK25 | *A. seifertii* UBA3035 | *A. seifertii* DSM102854 (NIPH 973) | *A. baumannii* ATCC17978 |
| AB348-IK22 | 100 (100) | 100 (100) | 100 (100) | 100 (100) | 97.1 (75.2) | 97.4 (75.2) | 90.26 (62.3) |
| AB359-IK23 | 100 (100) | 100 (100) | 100 (100) | 100 (100) | 97.1 (76.6) | 97.4 (75.4) | 90.26 (62.8) |
| AB349-IK24 | 100 (100) | 100 (100) | 100 (100) | 100 (100) | 97.1 (76.1) | 97.4 (75.2) | 90.26 (63.8) |
| AB350-IK25 | 100 (100) | 100 (100) | 100 (100) | 100 (100) | 97.1 (77.5) | 97.4 (75.1) | 90.28 (63.0) |
| *A. seifertii* UBA3035 | 97.1 (75.2) | 97.1 (76.6) | 97.1 (76.1) | 97.1 (77.5) | 100 (100) | 97.14 (82.3) | 90.26 (63.8) |
| *A. seifertii* DSM102854 (NIPH 973) | 97.4 (75.2) | 97.4 (75.4) | 97.4 (75.2) | 97.4 (77.5) | 97.14 (82.3) | 100 (100) | 90.36 (62.9) |
| *A. baumannii* ATCC17978 | 90.26 (62.3) | 90.26 (62.8) | 90.26 (63.8) | 90.28 (63.0) | 90.26 (63.8) | 90.36 (62.9) | 100 (100) |

However, AB348-IK22 has a dDDH value of 62.3 % compared to *A. baumannii* ATCC17978. All *A. seifertii* tank milk strains have a comparative dDDH value below the 75% threshold compared to *A. baumannii* and above this threshold compared to the two *A. seifertii* type strains (31). With high confidence based on these values, our tank milk strains AB348-IK22, AB359-IK23, AB349-IK24 and AB350-IK25 are identified as *A. seifertii*.

Epidemiological tracking of members of the ACB complex can be completed in a number of ways, but the most common is by Multi-Locus Sequence Typing (MLST). According to the Pasteur MLST scheme, AB348-IK22, AB359-IK23, AB349-IK24 and AB350-IK25 have been assigned to the novel Sequence Type (ST), ST-2706 (data not shown)

### *Acinetobacter seifertii* strains are resistant to colistin

To determine their antibiotic susceptibility profiles, these strains were subjected to broth microdilution testing to the Canadian ward (CANWARD) (32) panel of antibiotics. There are no clinical breakpoints for *A. seifertii* specifically, but CLSI breakpoints are available for *Acinetobacter* spp. and were applied to *A. seifertii*. AB350-IK25 is resistant to ceftazidime (CAZ) and colistin (CST) and has an intermediate susceptibility to ceftriaxone (CRO). AB348-IK22, AB348-IK24 and AB359-IK23 show intermediate susceptibility to CAZ, CRO and also demonstrates resistance to CST The type-strain DSM102854 is susceptible to CST but shows intermediate resistance to CAZ, while *A. baumannii* type strain ATCC17978 is susceptible to all antibiotics tested except for intermediate susceptibility to CRO (Table 2).

**Table 2:** Antibiotic susceptibility of *A. seifertii*. Susceptibility testing results for the CANWARD panel of antibiotics according to the Clinical Laboratory Standards Institute (CLSI) broth microdilution guidelines. The Minimum Inhibitory Concentration (MIC) is shown in µg/mL. All values that indicate resistance are highlighted in bold text and those that indicate intermediate resistance are underlined. Data is displayed based on three biological replicates.

| Strain | CFZ | FEP | FOX | CAZ | BPR | C/T | CRO | CIP | CLR | CLI | CST | DAP | DOX | ETP | GEN | IPM | LZD | MEM | NIT | TZP | TOB | SXT |
| --- | --- | --- | --- | --- | --- | --- | --- | --- | --- | --- | --- | --- | --- | --- | --- | --- | --- | --- | --- | --- | --- | --- |
| ATCC17978 | >128 | 4 | >32 | 8 | 0.5 | 2 | <u>16</u> | 0.25 | 32 | >8 | 2 | >16 | 0.25 | 4 | 1 | 0.25 | >16 | 0.5 | >512 | 4 | 1 | 4 |
| DSM102854 | >128 | 4 | >32 | <u>16</u> | 1 | 0.5 | 16 | 0.12 | 8 | >8 | 2 | >16 | 0.25 | 16 | <0.5 | 0.25 | >16 | 1 | 512 | 2 | <0.5 | ≤0.12 |
| AB348-IK22 | >128 | 4 | >32 | <u>16</u> | 0.5 | 1 | <u>32</u> | 0.25 | 8 | >8 | **8** | >16 | ≤0.12 | 4 | 2 | 0.25 | >16 | 0.25 | 256 | 8 | 1 | ≤0.12 |
| AB349-IK23 | >128 | 2 | >32 | <u>16</u> | 0.5 | 2 | <u>16</u> | 0.25 | 16 | >8 | **16** | >16 | ≤0.12 | 8 | 2 | 0.5 | >16 | 0.5 | 128 | 4 | 1 | ≤0.12 |
| AB350-IK25 | >128 | 4 | >32 | **32** | 1 | 4 | <u>16</u> | 0.5 | 16 | >8 | **8** | >16 | 0.25 | 8 | 4 | 0.25 | >16 | 0.5 | 512 | 8 | 1 | ≤0.12 |
| AB359-IK24 | >128 | 4 | >32 | <u>16</u> | 0.5 | 2 | <u>16</u> | 0.5 | 8 | >8 | **16** | >16 | 0.25 | 4 | 2 | 0.25 | >16 | 0.25 | 256 | 8 | 1 | ≤0.12 |

457 AMK; Amikacin, CFZ; Cefazolin, FEP; Cefepime, FOX; Cefoxitin, CAZ; Ceftazidime, BPR; Ceftobiprole, C/T; Ceftolazane/tazobactam, CRO; Ceftriaxone, CIP; Ciprofloxacin, CLR; Clarithromycin, CLI; Clindamycin, CST; Colistin, DAP; Daptomycin, DOX; Doxycycline, ETP; Ertapenem, GEN; Gentamicin, IPM; 458 Imipenem, LZD; Linezolid, MEM; Meropenem, NIT; Nitrofurantoin, TZP; Piperacillin/Tazobactam, TOB; Tobramycin, SXT; Trimethoprim/Sulphamethoxazole, VAN; Vancomycin

### The mechanism of colistin resistance in *A. seifertii* is distinct from known mechanisms in *A. baumannii*

Upon genomic investigation into the cause of this phenotype, no known colistin resistance mechanisms were detected when using the Resistance Gene Identifier with perfect and strict hit parameters from the Comprehensive Antibiotic Resistance Database (CARD). When including the loose hits as well as the perfect and strict (Figure 1), there are similar gene identities between colistin susceptible and non-susceptible strains. These include *bacA*, *kpnH, rosAB, tolC, lptD, ugd, pmrF, cpxR,* and *cls1.* Conservation of *lpsB* is present; ATCC17978 (NZ_CP018664.1) with 98.90% identity, DSM102854 with 95.63% identity and AB348-IK22, AB359-IK23, AB349-IK34 and AB350-IK25 with 95.08% identity. Sequences of the phosphoethanolamine transferase, *eptA,* were detected in the colistin-resistant *A. seifertii* strains including the type strain, DSM102854 (40.78% identity) and in colistin susceptible *A. baumannii* ATCC17978 (42.16% identity). The two-component system, *cprRS* is not detectable in ATCC17978 but appears to be present in all *A. seifertii* strains (34.55% for *cprR* and 26.43% for *cprS*). Genes from the *arn* operon, namely, *arnAT*, were detected in ATCC17978 (*arnA* at 27.91% identity and *arnT* at 31.15% identity) and in DSM102854 (*arnA at* 28% identity and *arnT* at 28.01% identity) and in AB348-IK22, AB359-IK23, AB349-IK24 and AB350-IK25 (*arnA* at 27.67% identity and *arnT* at 27.91% identity). One detectable difference between *A. seifertii* DSM102854 and the tank milk *A. seifertii* strains is the lack of the Resistance-Nodulation-Division (RND) efflux pump, *adeAB* and its cognate two-component regulatory system, *adeRS* (Figure 1). These four genes are undetectable in DSM102854. Since *adeAB* and its regulators, *adeRS,* are the only *Acinetobacter* spp. specific genes,, expression of *adeB* (as a proxy for the entire operon) was investigated. There was a significant down-regulation of *adeB* expression in AB348-IK22, AB359-IK23, AB349-IK24 and AB350-IK25 compared to *A. baumannii* ATCC17978 (Supplemental Figure 1).

**Figure 1:**
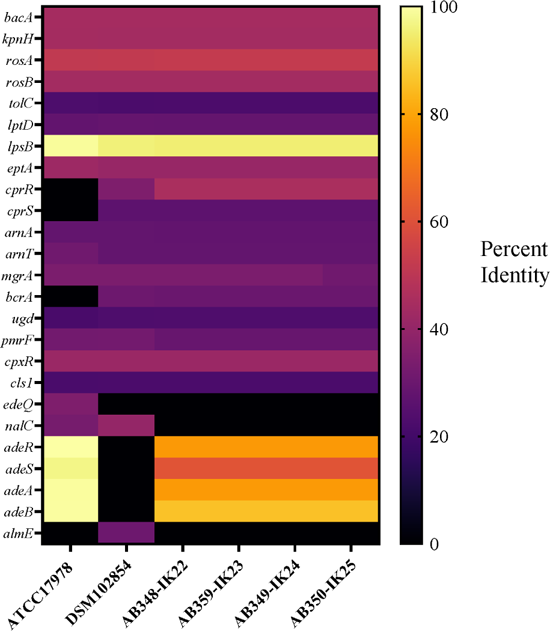
Colistin resistome for *A. seifertii* strains. Using the Resistance Gene Identifier (RGI) in the Comprehensive Antibiotic Resistance Database (CARD), colistin resistance genes were evaluated using perfect, strict and loose parameters. Genes associated with colistin resistance from the CARD are shown with the lighter the colour representing higher identity to that allele in the CARD. Black indicates that there was no gene in the genome that matched the one in the CARD.

Many of the genes detected in the CARD analysis are not specific to *Acinetobacter* spp. (Figure 1). Thereby, further investigation was done via a specific *A. baumannii* literature search on colistin resistance, using tblastn (33) to search for sequences manually in all genomes (Supplemental Table 1). This search yielded no further information regarding characterized insertional elements or mutations that may confer resistance to colistin in *A. seiferttii*. There are, however, amino acid substitutions in LpxDCA, PmrAB, H-NS, MlaD, PldA, AdeRS, LpsB and EptA (Supplemental Figures 3-10), all of which are uncharacterized with regards to colistin resistance, except in AdeS (V27L and L172P) (Supplemental Table 1 and Supplemental Figure 8b). Further investigation into the impact of these substitutions is required. It is noticeable that LpxC in colistin-resistant AB348-IK22, AB359-IK23, AB349-IK24 and AB359-IK25 shows a S147R mutation compared to colistin susceptible *A. baumannii* ATCC17978 and a Q147R mutation compared to colistin susceptible *A. seifertii* DSM102854 (Supplemental Figure 5B).

Further, in LpxA, S237I is detected in AB348-IK22, AB359-IK23, AB349-IK24 and AB359-IK25 compared to both ATCC17978 and DSM102854 (Supplemental 5C). In PldA (Supplemental Figure 7), mutations K82E (compared to ATCC17978), Q82E (compared to DSM102854), L185W (compared to both ATCC17978 and DSM102854) D290T (compared to ATCC17978) and A290T (compared to DSM102854) were observed.

### Lipidomics identified lyso-phosphatidylethanolamine and an acyl homoserine lactone in higher abundance in *A. seifertii* than in *A. baumannii*

Total lipids were extracted and analyzed via LC-MS to investigate possible membrane variance that might account for this colistin resistance phenotype. The global lipidome response for each strain was visualized using Principal Component Analysis (PCA) and was used to evaluate outliers in the data (Supplemental Figure 11). The colistin-sensitive *A. baumannii* ATCC17978, clustered tightly with *A. seifertii* strain AB348-IK22 (Supplemental Figure 11, pink and red data points). AB349-IK24 and AB350-IK25 cluster separately from other strains (Supplemental Figure 11, green data and blue data respectively). Colistin-sensitive *A. seifertii* DSM102854 (yellow data in Supplemental Figure 11) clusters most closely with AB359-IK23 (cyan data in Supplemental Figure 11).

A total of 559 compounds were identified (Supplemental Table 2) with 30 of them being significantly differential compared to ATCC17978 (p≤0.0001) (Figure 2). There was no detection of phosphoethanolamine modification in the significantly different compounds nor in the remaining results. Metabolic enrichment analysis based on KEGG pathways reveals that the most highly enriched lipids are involved in the biosynthesis of unsaturated fatty acids, synthesis and degradation of fatty acids as well as butanoate metabolism (Supplemental Figure 12). The least enriched lipids were in fatty acid elongation, bile acid metabolism and steroid hormone biosynthesis.

**Figure 2:**
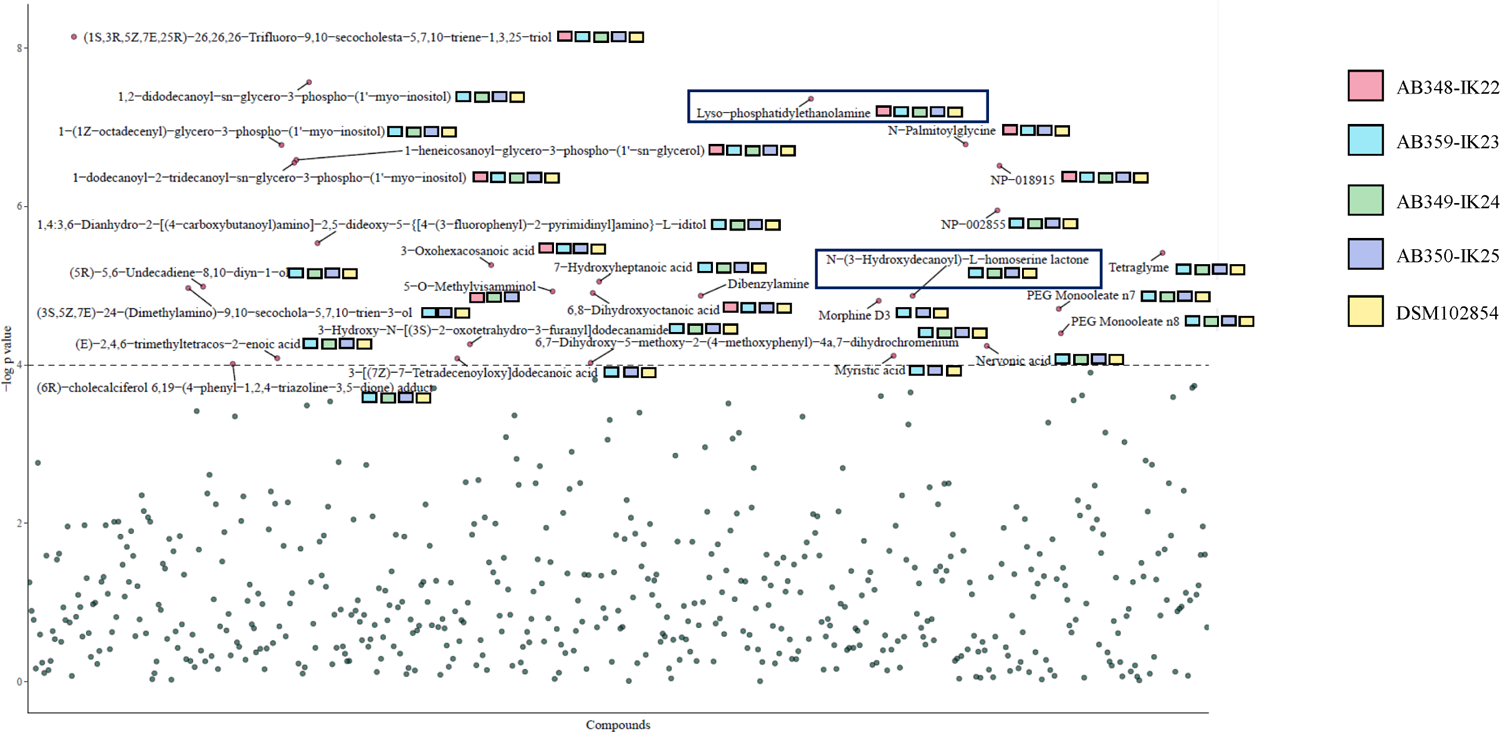
Global lipid profile of all *A. seifertii* strains compared to *A. baumannii* ATCC17978. All compounds are shown with those of statistical significance indicated in pink and above the dotted line (p (p≤0.0001). Each square next to the compound name indicates which *A. seifertii* lipid sample contained a greater abundance than *A. baumannii* ATCC17978. Lyo-phoshpatidylethanolamine (Lyso-PE) and N-(3-Hydroxydecanoyl)-L-homoserine lactone are highlighted with dark blue boxes to indicate their importance.

The top differential compounds revealed two interesting differences: lyso-phosphatidylethanolamine (lyso-PE) and N-(3-hydroxy-dodecanoyl)-homoserine lactone (Figure 2). Lyso-PE is a known component of *A. baumannii* membranes and a higher abundance of lyso-glycerophospholipids has been linked to membrane remodelling due to loss of LOS and thus colistin resistance (34). However, the LOS has not been lost in the case of these *A. seifertii* strains (Supplemental Figure 2). The banding pattern of the extracted LOS is the comparable between *A. baumannii* ATCC17978, and the *A. seifertii* strains AB348-IK22, AB359-IK23, AB349-IK24 and AB350-IK25.

In Figure 2, N-(3-hydroxy-dodecanoyl)-homoserine lactone (OH-C10-DHL) was detected at a significant level above ATCC17978 in strains AB359-IK23 (cyan box), AB349-IK24 (green box), AB350-IK25 (blue box) and DSM102854 (yellow box). OH-C10-DHL is a member of the acyl homoserine lactone family of compounds and is a known quorum sensing (QS) molecule in *A. baumannii* (35). Homologs to the sensor kinase of the LuxPQ quorum sensing system were identified in all six genomes and *rhtB* (annotated as homoserine lactone efflux protein) homologs were detected in DSM102854, AB348-IK22, AB359-IK23, AB349-IK24 and AB350-IK25 (data not shown).

Detection of the AHL in the lipidomics data led an investigation into QS-mediated phenotypes such as biofilm formation and motility. AB348-IK22 and AB359-IK23 showed significantly higher biofilm formation than ATCC17978 (Figure 4a), while AB350-IK25 was the only strain less motile than ATCC17978 (Figure 4b).

### *A. seifertii* tank milk strains are avirulent compared to *A. baumannii*

Virulence is an important characteristic to understand in any pathogen as it refers to its ability to cause infection. In *Acinetobacter* research, the *Galleria mellonella* wax larvae model has been optimized (36). Additionally, due to the fact that most *Acinetobacter* spp. research has been completed in *A. baumannii* only ATCC17978 was included for comparison. Over the total time of study (72 hours), there was a significantly higher survival rate of the *G. mellonella* larvae for all *A. seifertii* tank milk strains (Figure 3). All of the larvae survived upon injection with these strains. AB348-IK22, AB359-IK23, AB349-IK24 and AB350-IK25 are statistically less virulent compared to *A. baumannii* ATCC17978.

**Figure 3:**
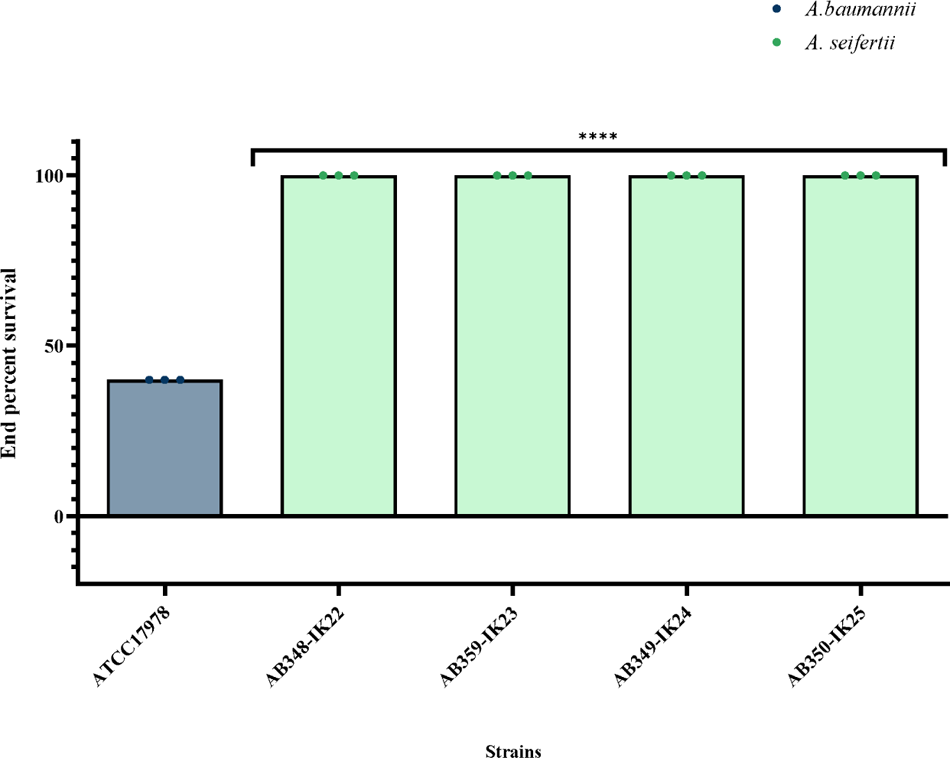
Virulence of *A. seifertii* tank milk strains in the insect model *Galleria mellonella.* Survival of injected larvae are counted over 72 hours at 37°C. One-Way ANOVA with Dunnett’s post-hoc analysis was used to determine significance. **** p < 0.00001. GraphPad Prism 10.0.0 was used for statistics.

**Figure 4:**
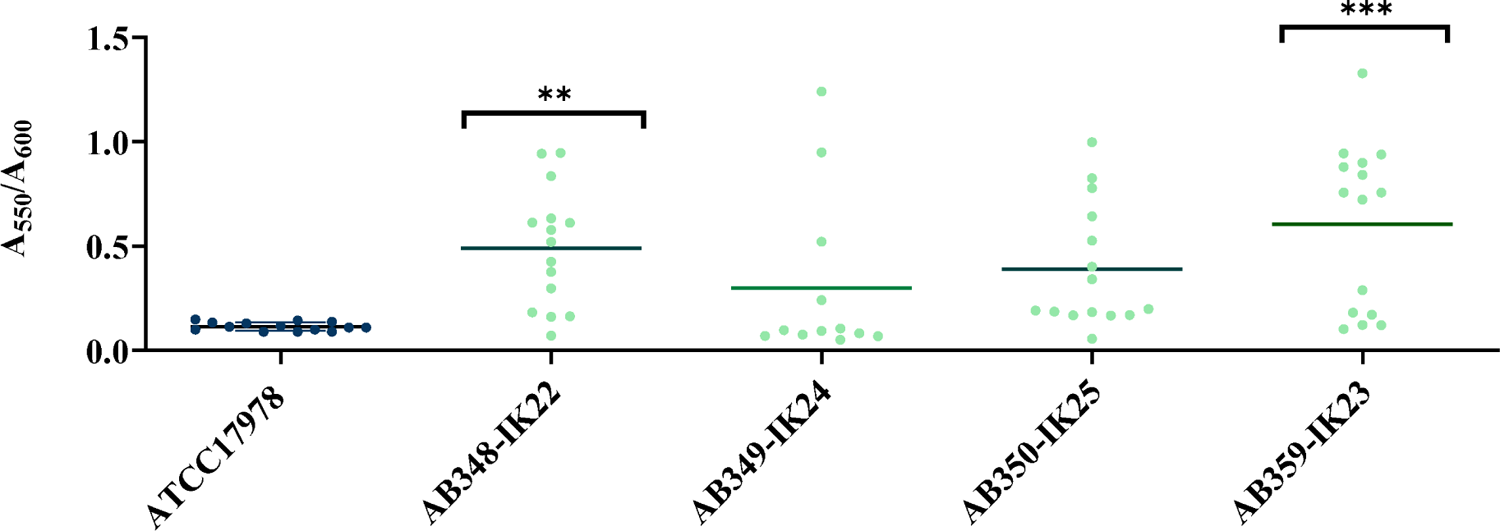
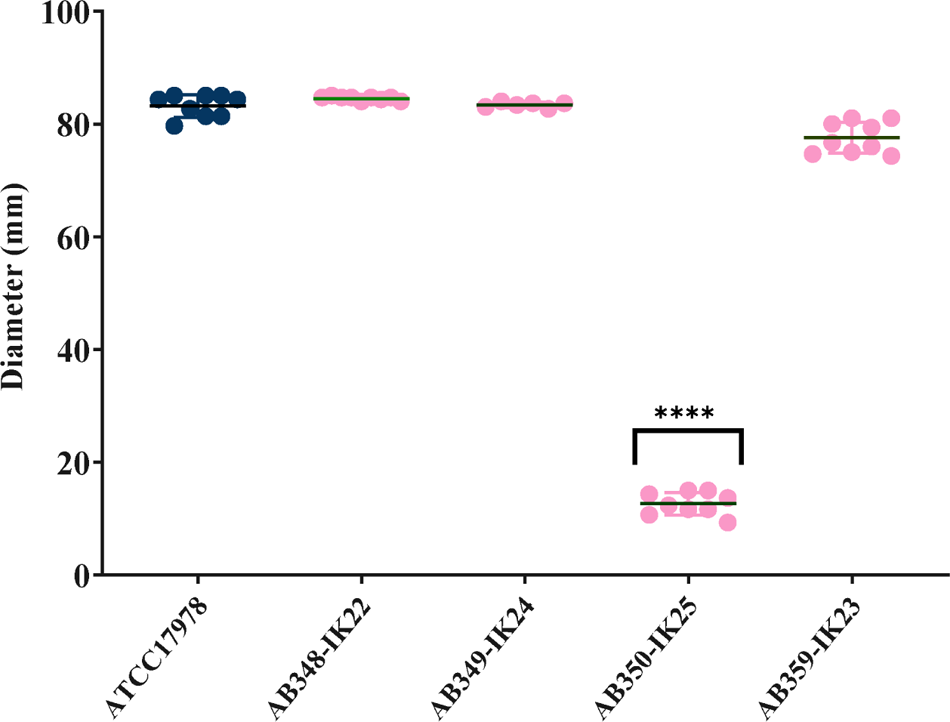
Quorum sensing-associated phenotypes in *A. seifertii*. **A:** Biofilm formation of *A. seifertii* strains. Biomass of biofilm was determined with crystal violet staining. Three biological replicates with four technical replicates were performed. One-way ANOVA with Dunnett’s post-hoc testing was used to determine statistical significance. **B:** Motility of *A. seifertii* strains. The distance travelled on semi-solid media was measured after 18 hours of incubation at 37°C. Three biological replicates with three technical replicates were performed. One-way ANOVA with Dunnett’s post-hoc testing was used to determine statistical significance.

## Discussion

All *A. seifertii* tank milk strains have a comparative dDDH value below the 75% threshold compared to *A. baumannii* and above this threshold compared to the two *A. seifertii* type strains. Colistin has been used mainly as a last-resort antibiotic in clinical settings but is also used in agricultural and veterinary practices as a means of infection prevention. The fact that non-clinical isolates of *A. seifertii* possess resistance to colistin is intriguing and contributes to the theory that *A. seifertii* is intrinsically resistant to colistin. Not much is known about the molecular resistance mechanisms of this ACB complex member. Upon investigation using the CARD to determine the ARGs responsible, known genetic determinants of colistin resistance were not revealed (Figure 1). We observed other genetic differences between *A. baumannii* ATCC17978, the *A. seifertii* type strain (DSM102854) and the four tank milk strains that may contribute to colistin resistance. These specific amino acid substitutions will be investigated in the future for their role in colistin resistance in *A. seifertii*. In a clinical isolate of *Acinetobacter colistiniresistens*, it was found that SNPs in *lpxACDL* could be responsible for its colistin-resistant phenotype (37). In fact, *A. colistiniresistens* is among only a few non-*baumannii Acinetobacter* spp. including *A. junii, Acinetobacter parvus, Acinetobacter beijerinckii*, and genomic species 16, recently identified as *A. higginsii* (38), to confer intrinsic resistance to colistin (39). SNPs were detected in the *lpxACDL* operon in our tank milk *A. seifertii* strains but none of them were common to *A. colisteniresistens* (Supplemental Figure 3-10). The variation in membrane biosynthetic genes may simply represent the evolution of membranes in *Acinetobacter* spp. The down regulation of *adeB* in AB348-IK22, AB359-IK23, AB349-IK24 and AB350-IK25 may also be contributing to the colistin resistance, however, it remains unclear as to the exact mechanism. Typically, expression of RND efflux pumps is positively associated with susceptibility; an increase in expression is linked to a decrease in susceptibility because antibiotics are substrates of efflux pumps. Both type strains *A. baumannii* ATCC17978 and *A. seifertii* DSM102854 are sensitive to colistin. The genes *adeAB* are absent in DSM102854 but exist in ATCC17978 and are typically induced upon exposure to substrates of this pump including aminoglycosides (40). An efflux-mediated colistin resistance mechanism in *A. baumannii* has been suggested. Yilmaz and colleagues (41) demonstrate that in a colistin-resistant strain, inhibition of efflux, using the efflux pump inhibitor carbonyl cyanide 3-chlorophenylhydrazone (CCCP), resulted in increased susceptibility to colistin. Various mutations in the two-component system, AdeRS, including V27L and L172P in AdeS were suggested to be responsible for the colistin-resistant phenotype (41). Both mutations are observed in colistin-resistant AB348-IK22, AB359-IK23, AB349-IK24 and AB350-IK25 and may explain the downregulation of *adeB* observed (Supplemental Figure 1). However, contrary to this, the AdeS mutations mentioned above are also detected in colistin-sensitive DSM102854. Elucidation of the link between *adeAB* expression and colistin resistance requires further study.

The global lipidomic analysis revealed many compounds with different production between strains. Our study is limited in that some compounds were not identified and are therefore a part of the so-called ‘dark metabolome’ (42). Additionally, identification was challenging because of the bias of databases towards eukaryotic lipids which limits the number of metabolites that can be accurately interpreted in the bacterial context. Alternate modifications have been characterized in other species such as the presence of ArnT, which add an aminoarabinose group to lipid A in *Bordetella bronchiseptica, Burkholderia* spp, *Klebsiella* spp., *E. coli, Shigella* spp., *Pseudomonas* spp., *Salmonella* spp., and *Yersinia* spp. (22). This has not been detected in *Acinetobacter* spp. to date and our study did not detect this modification either.

This is the first report of OH-C10-HSL being identified in *A. seifertii* (Figure 2). This molecule has been characterized in *A. nosocomialis* (43), and most significantly in *A. baumannii* (35). In fact, there are LuxQ homologues in our four *A. seifertii* strains (data not shown) further supporting the capability of *A. seifertii* to use QS as a means for regulating population activities. This is also the first report to detect lyso-PE as a component of the *A. seifertii* membrane. In contrast to *A. baumannii*, the detection of lyso-PE in our study, indicates that this glycerophospholipid is found in higher abundance.

Despite the colistin-sensitive nature of *A. seifertii* type strain DSM102854, there are significant similarities between it and the colistin-resistant tank milk *A. seifertii* isolates. All of those compounds detected in the tank milk strains in higher abundance compared to *A. baumannii* ATCC17978, were also present in DSM102854 (yellow boxes, Figure 2). This suggests that there is some fine-tuned control of other processes within the cell that were not detectable via our methods. For example, there are no significant antibiotic resistance gene differences between the strains, apart from the high percent identity of *A. baumannii* RND efflux pump *adeAB* (Figure 1). The low expression of *adeAB* in AB348-IK22, AB359-IK23, AB349-IK24 and AB350-IK25 may be contributing to their colistin-resistant phenotype. Investigation into the global expression changes in these isolates may explain the link between little to no lipidomic differences between colistin-resistant and colistin-susceptible *A. seifertii*.

There is much information to be gained from the study of non-human ACB complex members. This study is one such example of the exploration of the diversity of this species, including the discovery of the novel ST status of these *A. seifertii* isolates. It also contributes to the importance of considering One Health in studying antimicrobial resistance.

## Methods

### Isolation and Culture Conditions

*A. seifertii* strains, AB348-IK22, AB359-IK23, AB349-IK24 and AB350-IK25 in this study were isolated from tank milk from Bogor, Indonesia, and grown using tryptic soy agar (5% sheep’s blood) and CHROMAgar *Acinetobacter* selective growth media at 30°C for 48 hours. They were isolated as a part of the collection in Sykes *et al* (12)*. A. seifertii* DSM102854 was purchased through the DSMZ culture library.

### Whole Genome Sequencing

Genomic DNA was extracted using the DNAeasy UltraClean microbial kit (Qiagen, MD, USA) from a purified colony of each strain according to the manufacturer’s instructions. Sequence libraries were prepared and pooled using the DNA prep and the NextSeq 500 mid-output reagent kits (Illumina, CA, USA). Illumina NextSeq 500 platform, at the Agriculture and Agri-Food Canada, Ottawa Research and Development Centre (AAFC-ORDC), was used for whole-genome sequencing, *de novo* genome assembly used SPAdes v. 3.12.0 (44). Completeness and quality of genome assemblies was performed using QUAST v 5.0.2 (45) and CheckM v1.0.11 (46) with a 95% completeness and equal or less than 5% contamination accepted. The sequences have been deposited to NCBI GenBank with the following: bioproject: PRJNA819071, biosample SAMN26898588 – SAMN26898591 and corresponding accession JANBNW000000000, JANBNX000000000, JANBNY000000000, JANBNZ000000000.

### Species Determination and MLST Analysis

The ANI based on the MUMmer alignment tool (ANIm) was used to determine the species. This was performed by uploading all assemblies AB348-IK22, AB359-IK23, AB349-IK24, AB350-IK25, *A. seifertii* UBA3035 (Accession: GCA_002367535.1), *A. seifertii* DSM102854 (NIPH 973) (Accession: GCF_000368065.1) and ATCC17978 (NZ_CP018664) to the Jspecies server (31). The calculation of dDDH was performed using The Type (Strain) Genome Server (47) by uploading all previously noted assemblies. MLST was performed using the Pasteur scheme (48) as reference via ABRicate (49). When no hits were observed, the novel ST profile were uploaded to pubMLST.org to await ST assignment. Genomes for the novel ST were uploaded as well. In order to determine Clonal Complex (CC) assignment, Phyloviz (50) was used to facilitate the goeBURST algorithm of analysis (51). CC assignment was determined based on those STs that share identical alleles at six of seven loci and the CC was named according to the founder ST. If no founder ST was determined, then that ST remains unassigned to a CC.

### MIC Determination

Antibiotic susceptibility was evaluated as per the CLSI guidelines using broth microdilution methods. The full CANWARD panel (32) of antibiotics was used for testing. Based on CLSI breakpoints (52), isolates were categorized into susceptible (S), intermediate (I) and resistant (R).

### Colistin Resistome Evaluation

All genome sequences were analyzed using the Resistance Gene Identifier (RGI) in the CARD (53) using perfect, strict and loose parameters. Percent identity was used to generate the heatmap in GraphPad Prism 10.0.0. Mutations associated with colistin resistance that have been characterized in *A. baumannii* were investigated in all *A. seifertii* strains.. Known sequences of genes involved in colistin resistance were compared using NCBI’s tblastn (54) against all genome sequences. Alignments were performed using MAFFT (55) and visualized using ESPript (56). Insertion sequences were evaluated using ISFinder (57) and any element that had an Evalue cutoff of >0.01 was considered a hit.

### Whole LOS Extraction and Staining

This protocol is adapted from Marolda *et al* (58). In brief, cultures are incubated in Luria Bertani (LB) broth (BD, New Jersey, USA) at 37°C for 18 hours with shaking. After pelleting the cells at an A_600_ of 1.5, lysis buffer is added, and the samples were boiled. DNase I and proteinase K were added at a final concentration of 100 µg/mL and 1.2 mg/mL respectively. After incubation with a hot phenol solution composed of 90% phenol, 0.1% 2-βmercaptoethanol, and 0.2% 8-hydroxyquinolone, the samples were chilled on ice and then centrifuged for phase separation.

The aqueous phase was removed and washed with ethyl ether. The LOS extracts were then run on NuPAGE Bis-Tris gradient (4-12%) SDS-PAGE gels (Invitrogen, Waltham, USA) with NuPAGE MES running buffer (Invitrogen, Waltham, USA) and stained with the Pierce Silver Stain kit (Invitrogen, Waltham, USA) according to the manufacturer’s protocol.

### Lipid A Extraction for Mass-spectrometry Analysis

Lipid A extraction was carried out using a previously published protocol (59) with a few modifications. In brief, 250 mL Lysogeny Broth (LB) (BD, New Jersey, USA) culture was inoculated and grown to A_600_ of 0.8 – 1.0. Cells were harvested and washed with 1 x phosphate-buffered saline (PBS). A single layer Bligh-Dyer reaction was then achieved by adding chloroform, methanol and water (1:2:0.8 v/v). After 20 minutes of incubation and centrifugation, an additional wash with the single-phase Bligh-Dyer mix was performed. Hydrolysis of LOS was performed with a mild acid buffer (50 mM sodium acetate pH 4.5, 1% SDS) and sonicated at a constant duty cycle for 20s at 50% output. The samples were then boiled, and the solution was converted to a two-phase Bligh-Dyer reaction by adding chloroform, methanol and water (2:2:1.8 v/v). The lower phase was extracted and then the upper phase was washed twice with a two-phase Bligh-Dyer solution. After pooling of the lower phases, the samples were rotary evaporated with a Buchi R-100 (Buchi, Newcastle, USA) with ARCTIC A25 Refrigerated Circulators cooling system (Thermo Fisher Scientific, Waltham, USA) and stored at −20°C prior to transport on dry ice to BioZone (University of Toronto, Toronto, Canada) for LC-MS analysis.

### Liquid Chromatography – Mass-spectroscopy analysis

Samples were resuspended in chloroform and transferred to a glass vial then dried under nitrogen. A 10-μl aliquot of each sample was run in a randomized order on a ZIC-pHILIC (polymeric hydrophilic interaction chromatography) column (SeQuant) or a ZIC-HILIC (hydrophilic interaction chromatography) column (SeQuant) coupled to an Orbitrap mass spectrometer (Thermo Scientific) or an Orbitrap Q Exactive mass spectrometer (Thermo Scientific) according to previously published methods (60). Both positive and negative modes were used with a full scan range m/z 80–1200, resolution 70,000 at 1 Hz, automatic gain control (AGC) target of 3 × 106, and a maximum injection time of 100 ms.

Fragmentation of pHILIC column-separated metabolites was performed in a data-dependent manner on the Q Exactive (Thermo Scientific) mass spectrometer, with the five most intense ions picked in a 1 m/z exclusion window and at a normalized collision energy of 30. All other conditions were the same as previously reported (60). Raw spectral files have been uploaded to MetaboLights (61) with the unique identifier MTBLS9659 (www.ebi.ac.uk/metabolights/MTBLS9659).

Statistical analysis was conducted in MetaboAnalyst v 5.0 (62). PCA was used to determine the similarity in biological replicates for each sample. AB348-IK22, AB349-IK24, AB350-IK25 and DSM102854. Metabolite identification was performed using LIPIDMAPS (63) as the reference database via ChemSpider. All detected lipids are listed in Supplemental Table 2. In MetaboAnalyst, a one-way ANOVA with Fisher’s post-hoc testing was used to determine significant differences in lipids compared to *A. baumannii* ATCC17978. Enrichment analysis was performed using the KEGG pathway categories.

### Virulence in Galleria mellonella

Protocol based on Sykes *et al* (64). In brief, larvae were sorted based on size and all those that were deemed equal were used. Overnight cultures of *A. baumannii* and *A. seifertii* were standardized to 0.5 MacFarland and then diluted 1/100 in 0.85% sterile saline. The ventral side of each larva was swabbed with 70% ethanol in preparation for injection. Then 10 µL of standardized diluted culture was injected into the left hind proleg. Larvae were then incubated in sterile plastic petri plates at 37°C for 72 hours. Survival was measured every 12 hours. At the beginning of each injection session, sterile 1x PBS was injected into 10 larvae as well as at the end to ensure syringe washes were adequate. Ten technical replicates and at least three biological replicates were performed for each strain. Percent end-point survival was calculated, and One-way ANOVA with Dunnett’s post-hoc test in GraphPad Prism 10.0.0 was used to evaluate significance compared to *A. baumannii* ATCC17978.

### RND Efflux Expression Evaluation

Expression analysis was performed according to Sykes *et al* (12). In brief, overnight cultures of each strain were subcultured 1/100 in LB broth with shaking at 37°C until mid exponential phase (A_600_ of 0.7±0.5). Cells were pelleted and supernatant removed before storing at −70°C overnight. RNA was extracted using the Invitrogen Purelink RNA extraction kit (Thermo Fisher, Waltham, USA), according to the manufacturer’s protocol. After DNase treatment using the Invitrogen Purelink DNase kit (Thermo Fisher, Waltham, USA), cDNA was synthesized using the Invitrogen VILO cDNA kit (Thermo Fisher, Waltham USA) from 1 µg DNase treated RNA. The newly synthesized cDNA was diluted 1/5 prior to analysis using RT-qPCR. Applied Biosciences (Beverly Hills, USA) SYBR select master mix was combined with the appropriate volume of sterile mQH_2_O and primers to a final concentration of 200 nM. Reaction was run on an Applied Biosciences (Beverly Hills, USA) StepOnePlus system. RT-qPCR for each strain was performed with three technical replicates and at least three biological replicates per target gene. The Pfaffl method of analysis was used to calculate the relative fold change expression compared to ATCC17978. One-way ANOVA using Dunnett’s post-hoc analysis was performed using GraphPad Prism 10.0.0.

### Biofilm Formation

This technique is based on Sykes *et al*.(12). In brief, overnight cultures of each strain were measured using A_600_. Cultures were standardized to 1.0 inoculated in flat bottom polystyrene plates. Incubated at 37°C for 48 hours followed by measurement of A_600_. Removal of planktonic cells was then followed by staining of the biofilm with 0.1% crystal violet. Solubilization of the stained biofilm was done using 30% acetic acid for 30 minutes after which A_550_ was determined. Five technical replicates were performed with at least three biological replicates for each strain. One-way ANOVA with Dunnett’s post-hoc analysis was used in GraphPad Prism 10.0.0 to determine significance. ** p< 0.01, ***p<0.0001.

### Motility

This protocol is based on Sykes *et al*. (12) In short, overnight cultures of each strain were measured at A_600_. Cultures were standardized to A_600_ of 1.0. Then 3 µL was stab inoculated into the centre of semi-solid minimal media and incubated at 37°C for 18 hours. Three measurements around the diameter of distance travelled were recorded. At least three biological replicates with three technical replicates for each strain were performed. One-way ANOVA with Dunnett’s post-hoc analysis was used in GraphPad Prism 10.0.0 to determine significance. **** p<0.00001.

## Acknowledgements

The authors would like to thank Drs. A. Motnenko and A. Hogan for their protocol and expertise in LOS extraction and staining. We would also like to thank Nancy Laing for antibiotic susceptibility testing.

